## Supplementary material for "Chromatin accessibility variation provides insights into missing regulation underlying immune-mediated diseases": Suppl Figs + Suppl Tables 5+7+10

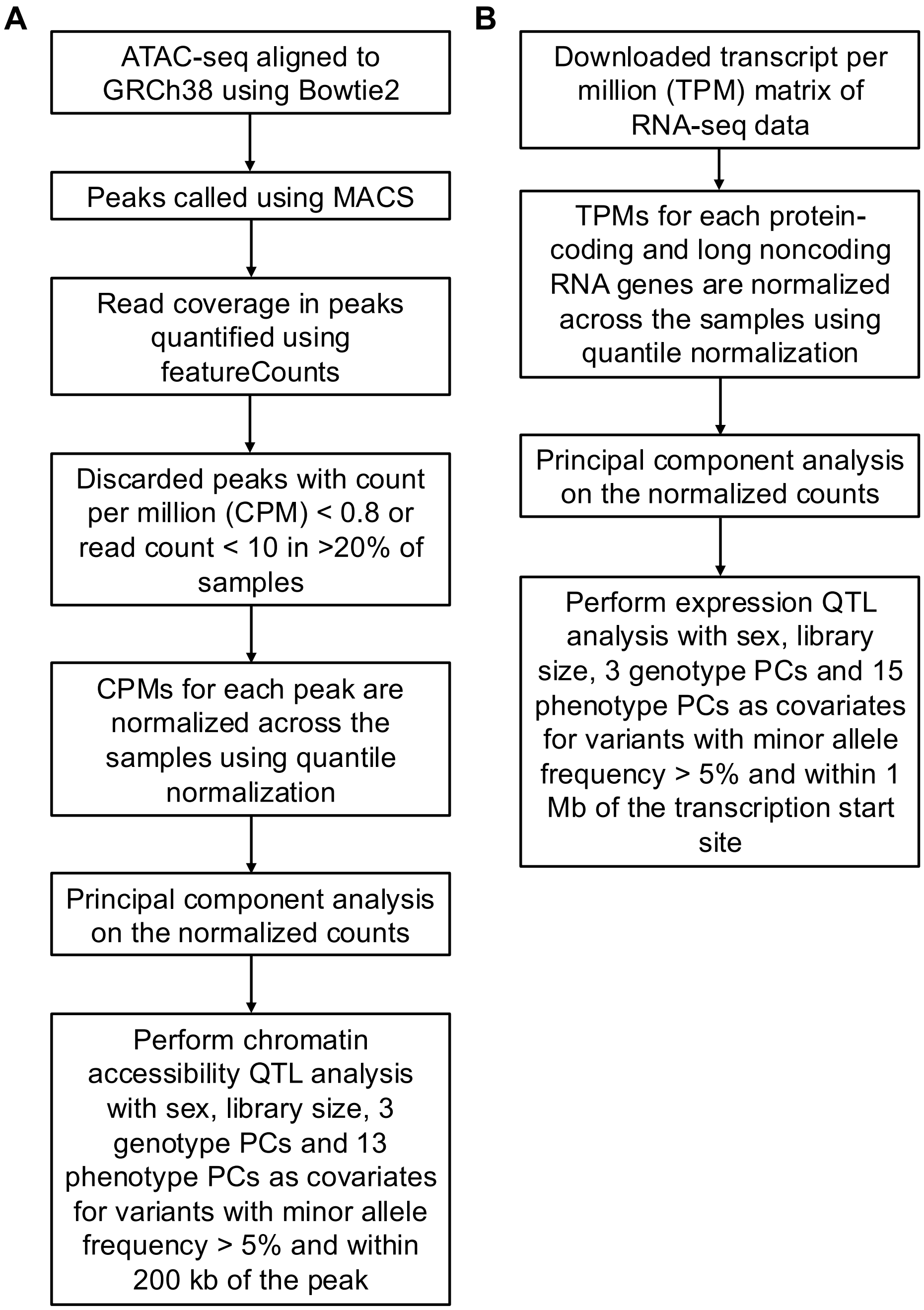


**Supplementary Figure 1. Workflow of caQTL and eQTL analyses.** (**A**) caQTL analysis workflow. (**B**) eQTL analysis workflow.

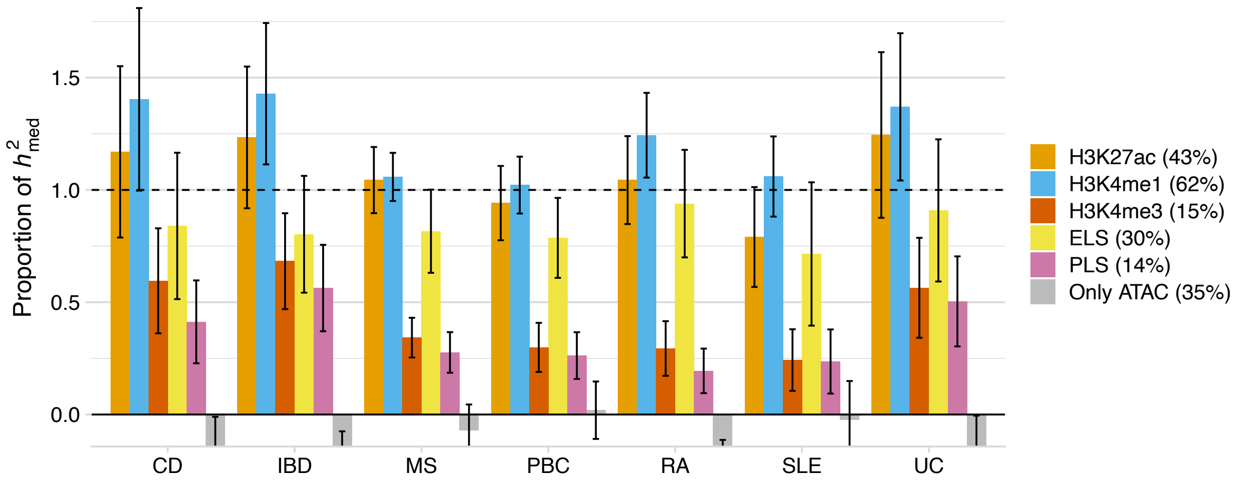


**Supplementary Figure 2. Proportion of caQTL-mediated IMD heritability explained by ATAC peaks with various histone marks.** 'Only ATAC' peak set includes peaks without any of the three histone marks (H3K27ac, H3K4me1, H3K4me3). Percentages for each peak set denote the proportion of ATAC peaks in that set. ELS, enhancer-like signature; PLS, promoter-like signature.

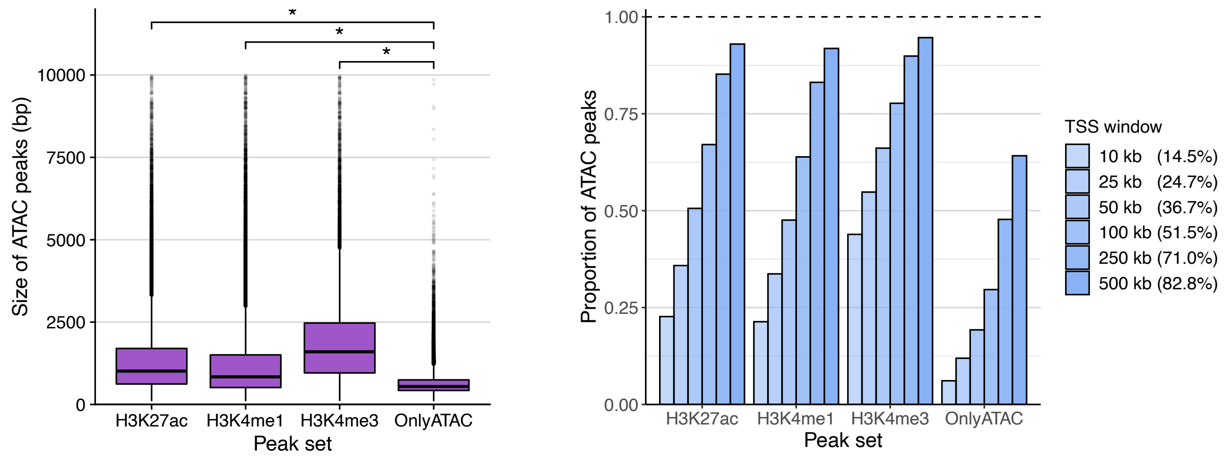


**Supplementary Figure 3. Properties of ATAC peaks with various histone marks.** (**A**) Size of ATAC peaks with various histone marks. Peaks with size greater than 10,000 bp are discarded. *: $p < 2.2\times10^{-16}$, Wilcoxon rank-sum test. (**B**) Proportion of ATAC peaks that are within the given TSS window around genes expressed in LCLs. Percentages for each TSS window denote the proportion of ATAC peaks in that window. 'Only ATAC' peak set includes peaks without any of the three histone marks (H3K27ac, H3K4me1, H3K4me3).

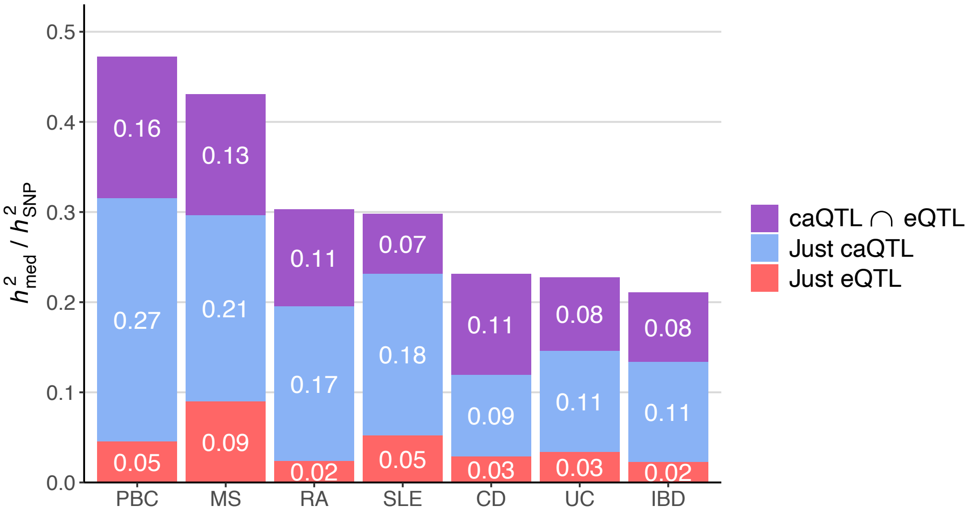


**Supplementary Figure 4. Relationship between IMD heritability mediated by caQTLs and eQTLs.** The numbers in the bars denote the estimated proportion of $h^2_{\mathrm{med}}$ / $h^2_{\mathrm{SNP}}$ by each subset. They are derived from $h^2_{\mathrm{med;\ caQTL}}$ / $h^2_{\mathrm{SNP}}$, $h^2_{\mathrm{med;\ eQTL}}$ / $h^2_{\mathrm{SNP}}$ and $h^2_{\mathrm{med;\ caQTL \cup eQTL}}$ / $h^2_{\mathrm{SNP}}$ as described in the Methods.

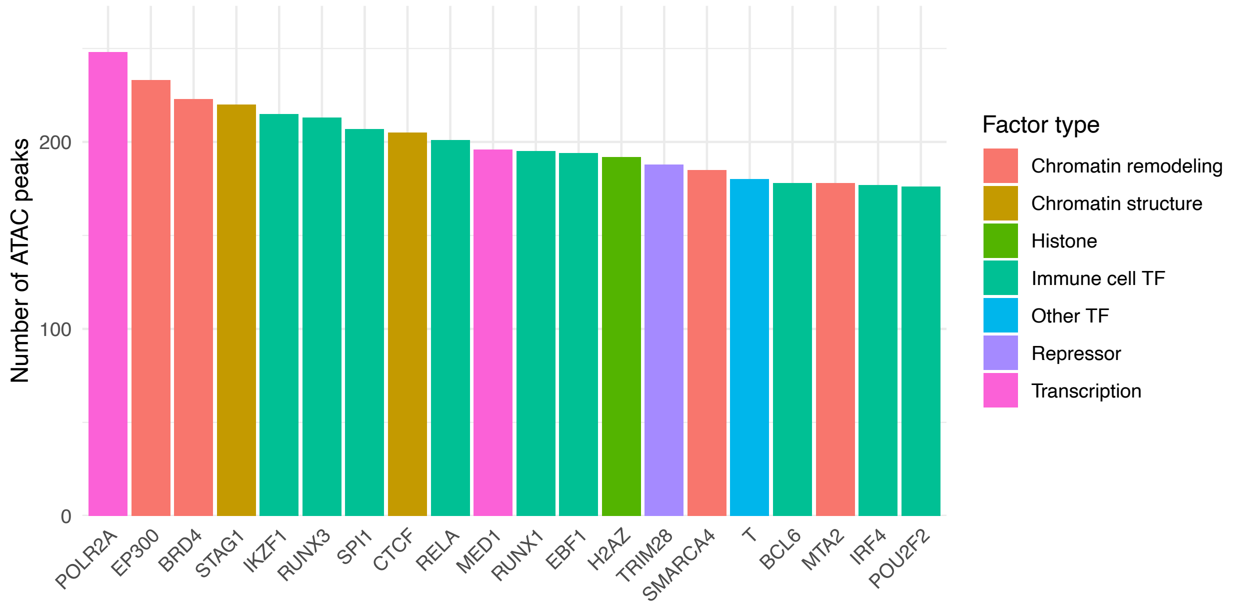


**Supplementary Figure 5. Protein factors detected at colocalized caQTL by ChIP-seq.** Number of unique ATAC-seq peaks overlapping ChIP-seq peaks of corresponding protein factors in Cistrome data. In total, there were 305 unique ATAC-seq peaks considered. Minimum overlap of 50% of the ChIP-seq peaks were counted. Only top 20 factors are shown.

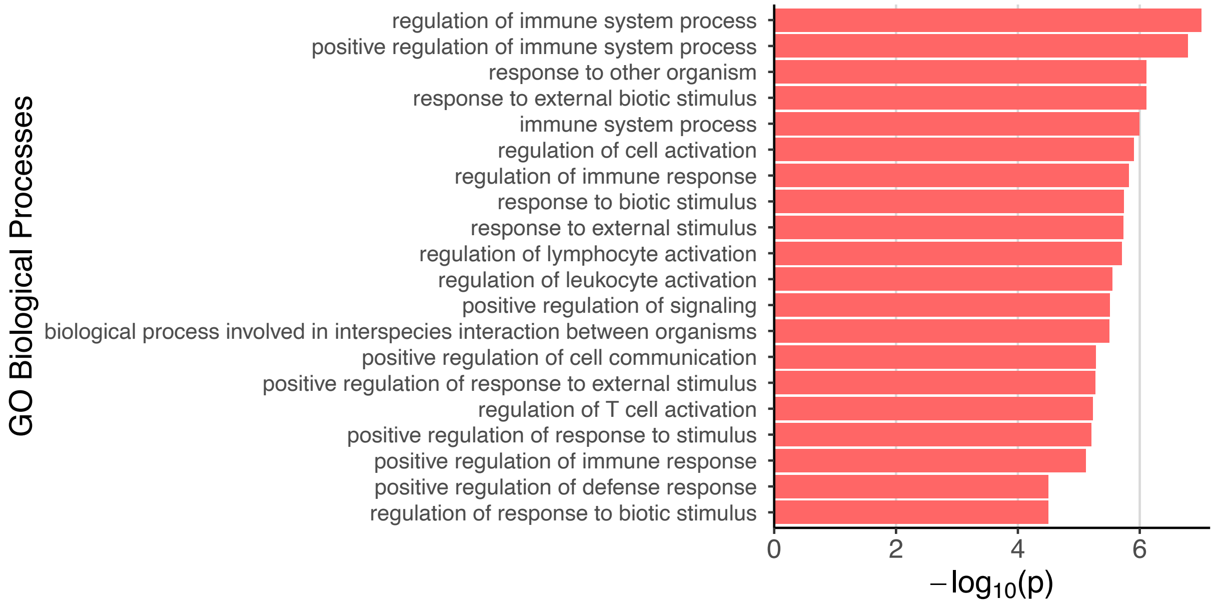


**Supplementary Figure 6. Biological processes enriched in IMD-colocalized eQTLs.** Top twenty enriched GO biological processes are shown. The p values were calculated using a binomial test in the Protein Analysis Through Evolutionary Relationships (PANTHER)[1] tool.

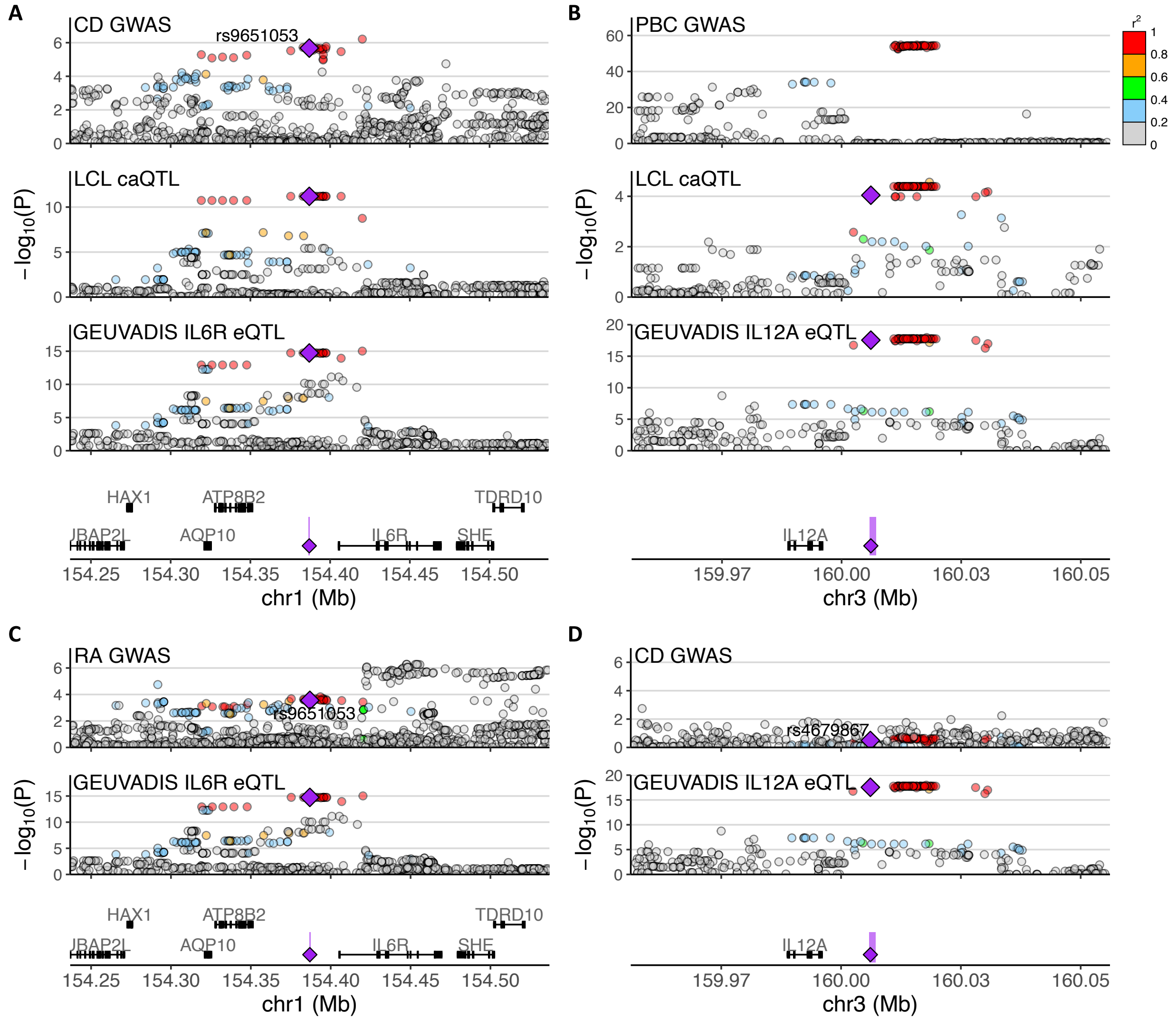


**Supplementary Figure 7. IMD GWAS colocalization with autoimmune disease drug target gene eQTL in LCLs.** (**A-D**) Association plots for IMD GWAS and LCL's caQTLs and *IL6R* (A and C) and *IL12A* (B and D) eQTLs. Panels A and B show significant eQTL colocalization to CD and PBC, respectively. Panels C and D show lack of colocalization to RA and CD, respectively, which are the diseases that the respective drugs treat. In each panel, the yellow shade signifies the caQTL peak, and the purple diamond shows a strongly associated variant that is within that peak. The other variants are colored by the degree of LD with the annotated variant. PBC GWAS data in (B) did not have the highlighted variant (rs4679867), so the purple diamond is missing.

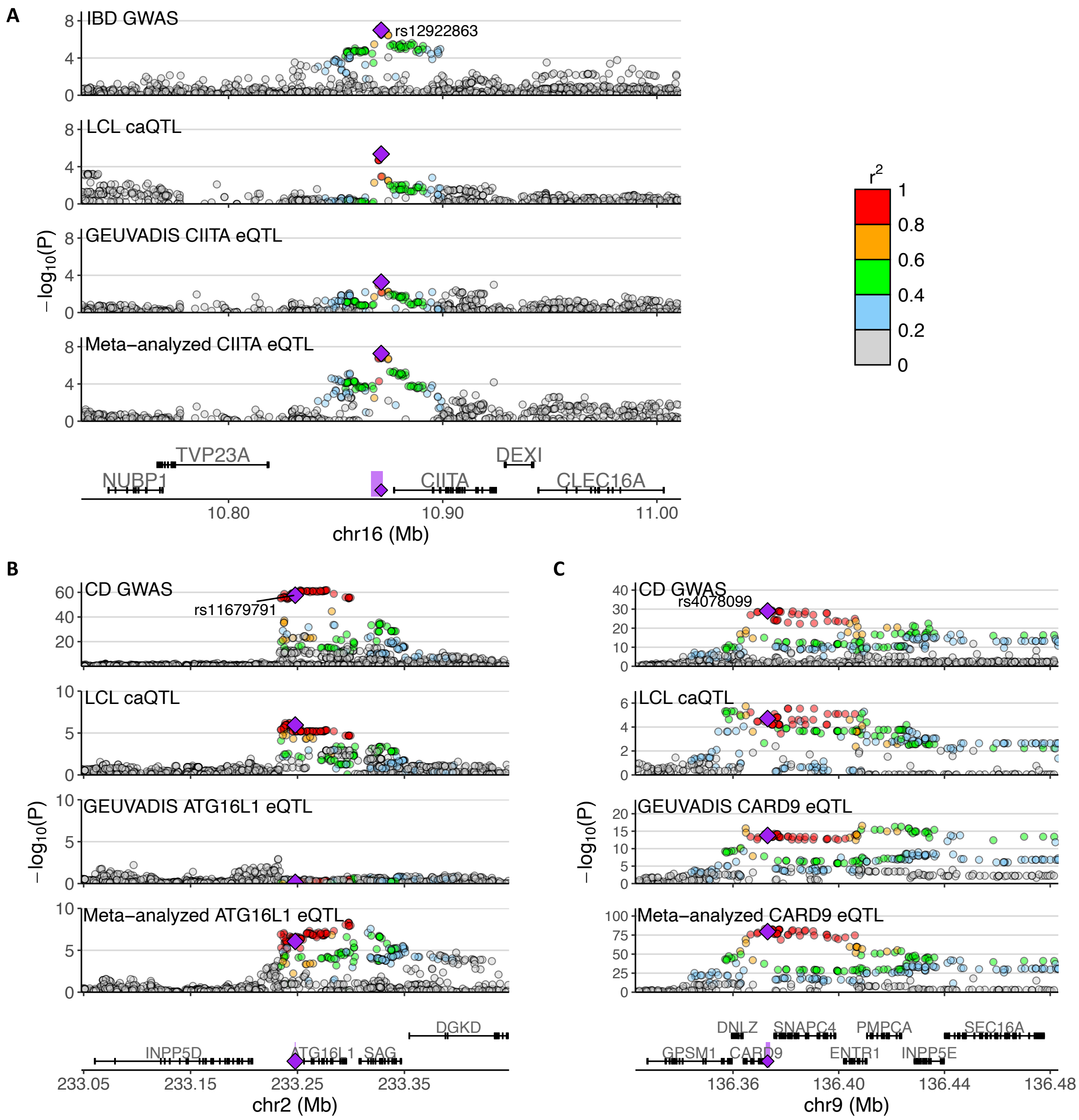


**Supplementary Figure 8. IMD GWAS colocalization with meta-analyzed LCL eQTLs.** (A-C) Association plots for IMD GWAS, caQTLs, and *CIITA* (A), *ATG16L1* (B), and *CARD9* (C) eQTLs. GEUVADIS eQTLs did not colocalize with the GWAS signals, but meta-analyzed eQTLs did. In each panel, the yellow shade signifies the caQTL peak, and the purple diamond shows a strongly associated variant that is within that peak. The other variants are colored by the degree of LD with the annotated variant.

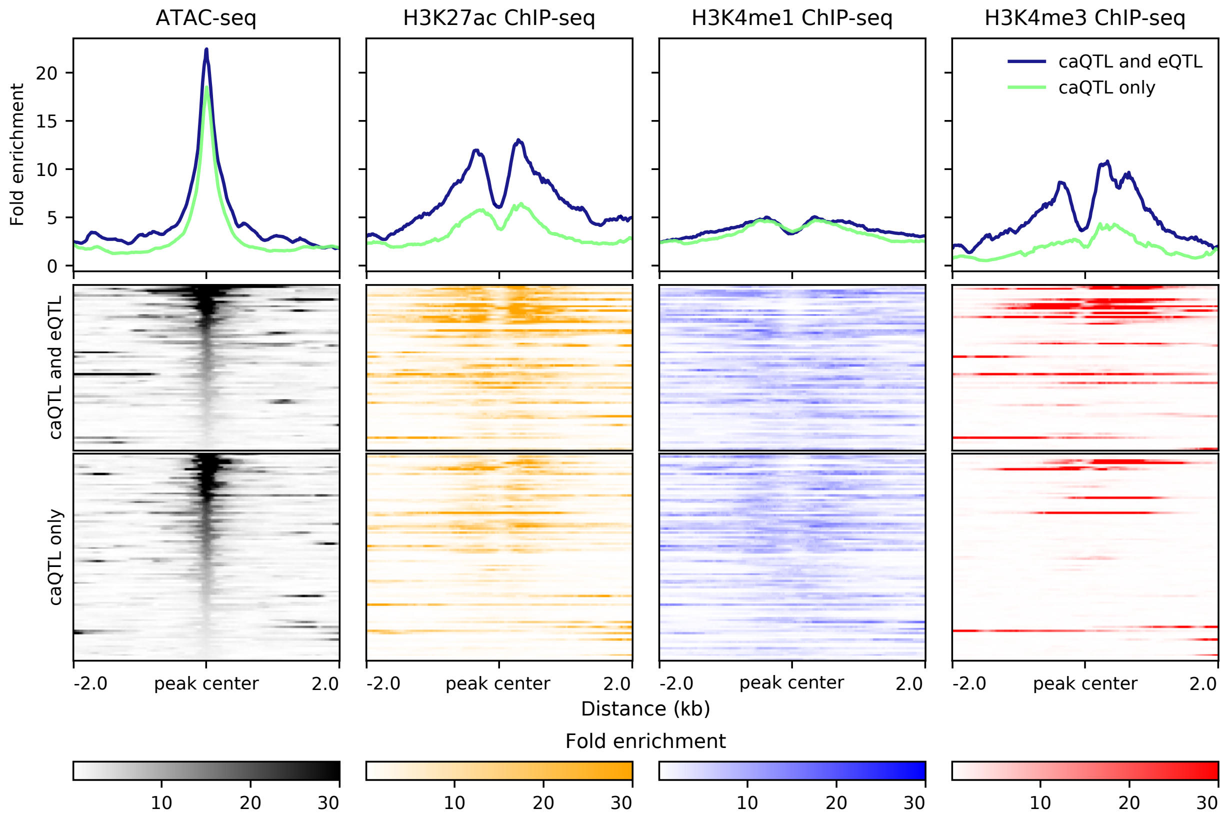


**Supplementary Figure 9. Chromatin accessibility and histone mark levels at IMD-associated LCL caQTLs.** IMD-associated caQTLs are separated by whether eQTL also colocalized with IMD association. (Top) Average profile of fold enrichment values of each assay in the 4-kb window centered at the caQTL peak center. (Bottom) Heatmap of fold enrichment values of each assay in the 4-kb window centered at the caQTL peak center.

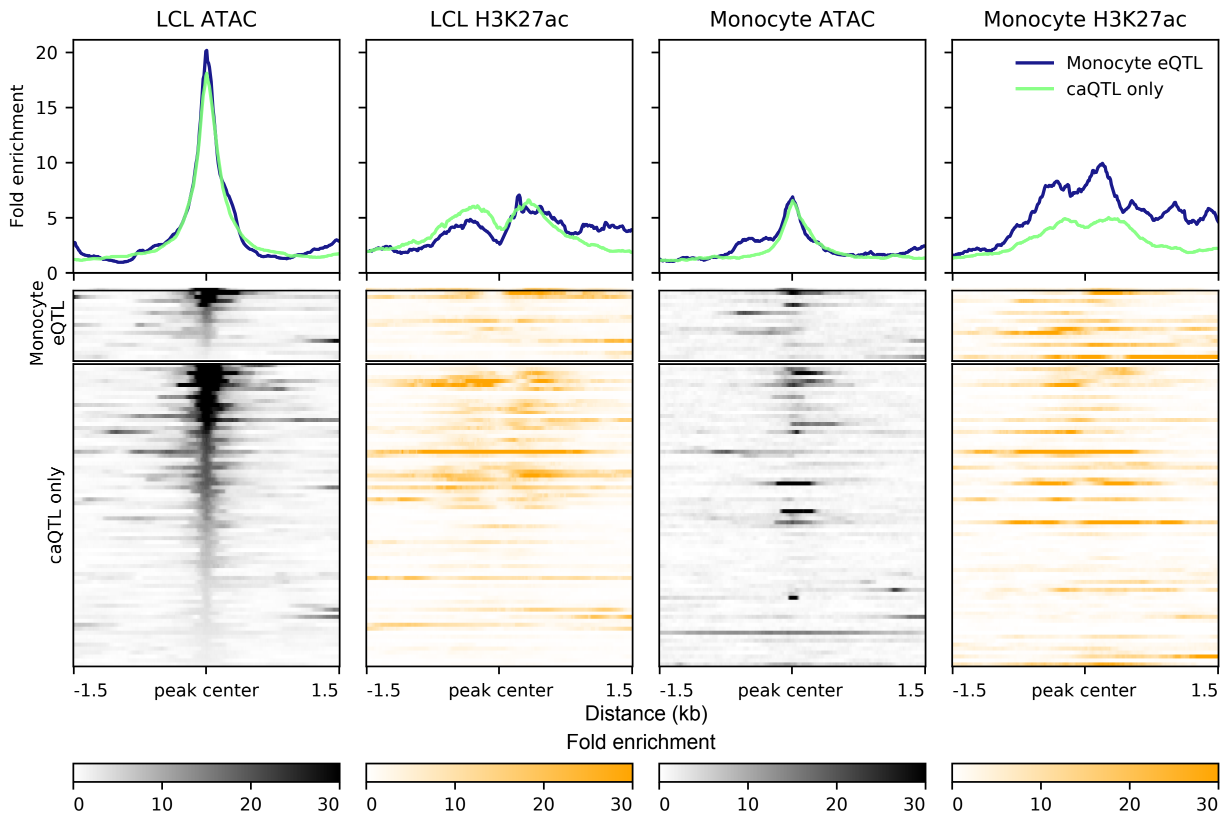


**Supplementary Figure 10. Chromatin accessibility and H3K27ac mark levels at IMD-associated caQTLs with respect to monocyte eQTL colocalization.** IMD-associated caQTLs without LCL eQTL colocalization are separated by whether monocyte eQTL colocalized with IMD association. (Top) Average profile of fold enrichment values of each assay in the 3-kb window centered at the caQTL peak center. (Bottom) Heatmap of fold enrichment values of each assay in the 3-kb window centered at the caQTL peak center.

**Supplementary Table 1 (See Excel file) Colocalization results of LCL caQTLs and IMD GWAS.**

**Supplementary Table 2 (See Excel file) Colocalization results of GEUVADIS eQTLs and IMD GWAS.**

**Supplementary Table 3 (See Excel file) Protein factor ChIP-seq peaks overlapping with colocalized caQTLs.**

**Supplementary Table 4 (See Excel file) Colocalization results of meta-analyzed LCL eQTLs and IMD GWAS.**

**Supplementary Table 5. Immune cell eQTL datasets analyzed in this study**

| Study | Cell type | Stimulated | Sample size |
|---|---|---|---|
| Schmiedel et al.[2] | Naïve B cell | naive | 91 |
| | Activated CD4$^{+}$ T cell | stimulated | 89 |
| | Naïve CD4$^{+}$ T cell | naive | 88 |
| | Activated CD8$^{+}$ T cell | stimulated | 88 |
| | Naïve CD8$^{+}$ T cell | naive | 89 |
| | CD16$^{+}$ monocyte | naive | 90 |
| | Classical monocyte | naive | 91 |
| | NK cell | naive | 90 |
| | $T_{FH}$ cell | naive | 89 |
| | $T_{H}$1/17 cell | naive | 88 |
| | $T_{H}$17 cell | naive | 89 |
| | $T_{H}$1 cell | naive | 82 |
| | $T_{H2}$ cell | naive | 89 |
| | Memory $T_{reg}$ cell | naive | 88 |
| | Naïve $T_{reg}$ cell | naive | 89 |
| BLUEPRINT[3] | CD4$^{+}$ T cell | naive | 167 |
| | Monocyte | naive | 191 |
| | Neutrophil | naive | 196 |
| Bossini-Castillo et al.[4] | $T_{reg}$ cell | naive | 119 |
| Alasoo et al.[5] | Naïve macrophage | naive | 84 |
| | Macrophage stimulated w/ IFNg | stimulated | 84 |
| | Macrophage stimulated w/ *Salmonella* | stimulated | 84 |
| | Macrophage stimulated w/ IFNg & *Salmonella* | stimulated | 84 |
| Soskic et al.[6] | Memory CD4$^{+}$ T cell | naive | 100 |
| | Memory CD4$^{+}$ T cell 16h | stimulated | 95 |
| | Memory CD4$^{+}$ T cell 40h | stimulated | 89 |
| | Memory CD4$^{+}$ T cell 5d | stimulated | 90 |
| | Naïve CD4$^{+}$ T cell | naive | 99 |
| | Naive CD4$^{+}$ T cell 16h | stimulated | 99 |
| | Naïve CD4$^{+}$ T cell 40h | stimulated | 89 |
| | Naïve CD4$^{+}$ T cell 5d | stimulated | 85 |

**Supplementary Table 6. (See Excel file) Colocalization results for immune cell eQTLs and IMD GWAS.**

**Supplementary Table 7. Overlap of LCL's caQTL and non-LCL eQTLs.**

| IMD | Total | caQTL & eQTL | caQTL only | eQTL only | no colocalization | Odds ratio | p value |
|---|---|---|---|---|---|---|---|
| CD | 196 | 27 | 12 | 64 | 93 | 3.2695 | $1.25\times10^{-3}$ |
| IBD | 259 | 34 | 14 | 74 | 137 | 4.4961 | $6.36\times10^{-6}$ |
| MS | 124 | 28 | 12 | 40 | 44 | 2.5667 | 0.0152 |
| PBC | 82 | 21 | 9 | 18 | 34 | 4.4075 | $1.98\times10^{-3}$ |
| RA | 146 | 25 | 11 | 44 | 66 | 3.4091 | $1.89\times10^{-3}$ |
| SLE | 72 | 6 | 6 | 18 | 42 | 2.3333 | 0.157 |
| UC | 142 | 18 | 7 | 42 | 75 | 4.5918 | $9.97\times10^{-4}$ |

**Supplementary Table 8. (See Excel file) Numbers of significant IMD-eQTL colocalization loci across IMDs and cell types.**

**Supplementary Table 9. (See Excel file) Colocalization results for meta-analyzed immune cell eQTLs and IMD GWAS.**

**Supplementary Table 10. 1000 Genomes samples in this study that required imputation.**

| Population | Sample ID |
|---|---|
| British in England and Scotland | HG00098 |
| | HG00104 |
| | HG00124 |
| | HG00134 |
| | HG00135 |
| | HG00152 |
| | HG00156 |
| | HG00247 |
| | HG00249 |
| Finnish in Finland | HG00312 |
| | HG00359 |
| | HG00377 |
| Toscani in Italy | NA20537 |
| | NA20816 |
